# DIFFERENTIAL EFFECTS OF PARAQUAT-INDUCED OXIDATIVE STRESS ON FUNCTIONAL AGING AND LIFESPAN IN MALE AND FEMALE DROSOPHILA MELANOGASTER

**DOI:** 10.1101/2025.10.22.684024

**Authors:** Mehmet Fidan

## Abstract

Aging is accompanied by loss of motor function driven by mitochondrial oxidative stress. To examine how genotype and sex modulate this process, we exposed Oregon-R (wild-type) and vestigial (wing-deficient) *Drosophila* of both sexes to chronic paraquat—a mitochondrial stressor generating sustained ROS production—at 0, 10, or 20 mM. We tracked climbing performance from day 5 to 50 alongside survival. Paraquat impaired locomotion dose-dependently, with effects modified by genotype and sex (four-way ANOVA, all p < 0.001). Under control conditions, behavioral half-life (T₅₀) occurred at 21.4 days in Oregon-R males and 25.7 days in females. Vestigial flies declined earlier: 14.8 days (males), 18.3 days (females). At 20 mM, T₅₀ fell 48-53% across groups. Female advantage persisted at 10 mM but narrowed at 20 mM, especially in vestigial flies. Survival mirrored functional decline. The T₅₀-to-lifespan interval compressed under severe stress: 18-28 days (controls) versus 8-12 days (20 mM). Functional-survival coupling was strong (r = 0.87, p < 0.001). Oxidative stress accelerates functional aging through mechanisms shaped by genotype and sex. Climbing performance predicts healthspan and may serve as a translational biomarker for neuromuscular aging.

## İntroduction

Aging is a gradual process that leads to the loss of physiological integrity and increased vulnerability to disease. This complex biological progression is characterized by nine interconnected hallmarks: telomere shortening, genomic instability, epigenetic modifications, mitochondrial dysfunction, loss of proteostasis, dysregulated nutrient sensing, stem cell exhaustion, cellular senescence, and altered cellular communication [1,2]. Among these mechanisms, oxidative stress has emerged as a central driver, linking multiple aging hallmarks through a common pathological pathway. According to the oxidative stress theory of aging, reactive oxygen species (ROS)—including superoxide radicals (O₂•⁻), hydrogen peroxide (H₂O₂), and hydroxyl radicals (•OH) are generated primarily through mitochondrial respiratory chain activity and endoplasmic reticulum stress responses [3]. These reactive species cause progressive damage to cellular macromolecules, including DNA oxidation, protein carbonylation, lipid peroxidation, and telomere shortening, ultimately leading to mitochondrial dysfunction, proteostasis collapse, and cellular senescence (Liguori et al., 2018; Moldogazieva et al., 2019). Importantly, chronic ROS accumulation dysregulates key metabolic signaling pathways, including Nuclear factor-κB (NF-κB), Mitogen-activated protein kinase (MAPK) cascades, Nuclear factor erythroid 2-related factor 2 (Nrf2), and phosphoinositide 3-kinase/protein kinase B (PI3K/Akt), which may trigger inappropriate cell death through apoptosis and necrosis [7,8]. The ability to manage such oxidative damage varies widely among individuals and has been linked to differences in longevity and healthspan [9].

The fruit fly *Drosophila melanogaster* is an established model for studying aging due to its short lifespan, well-characterized genetics, and conserved metabolic pathways. Its simple motor assays, particularly the negative geotaxis test, provide a sensitive readout of neuromuscular aging [10,11]. With recent advances in automated tracking systems, climbing behavior can now be quantified more precisely and reproducibly [12,13]. Because motor decline reflects both neuronal and muscular deterioration, this behavioral measure offers a useful indicator of overall healthspan.

Paraquat (1,1′-dimethyl-4,4′-bipyridinium dichloride) is a redox-cycling herbicide that serves as a powerful model for studying oxidative stress-induced neurodegeneration. At the molecular level, paraquat specifically targets mitochondria, where it undergoes single-electron reduction by Complex I of the electron transport chain to form paraquat cation radicals (PQ•⁺). These radicals rapidly react with molecular oxygen to regenerate paraquat and produce superoxide anions (O₂•⁻), creating a futile redox cycle that depletes NADPH and generates sustained oxidative stress [14,15]. This mechanism results in multiple mitochondrial defects, including decreased mitochondrial membrane potential (ΔΨm), reduced activities of respiratory chain complexes I-IV, impaired ATP synthesis, and increased mitochondrial fragmentation [15]. Beyond direct ROS generation, chronic paraquat exposure triggers secondary pathological cascades: lipid peroxidation produces reactive aldehydes such as 4-hydroxynonenal (4-HNE) and malondialdehyde (MDA), which form advanced lipoxidation end products (ALEs) that cause protein cross-linking and aggregation, leading to proteostasis collapse [4,16]. In *Drosophila*, these molecular events manifest as dopaminergic neuron loss, neuromuscular junction degeneration, and progressive locomotor impairment—hallmarks of Parkinson-like neurodegeneration [17–19]. Importantly, paraquat-induced oxidative stress also triggers neuroinflammation: studies in rodent models demonstrate that chronic oxidative stress elevates proinflammatory cytokines (IL-6, IL-1β) and neuroinflammatory markers (CD68, TLR4, MCP1), contributing to accelerated cognitive decline and functional deterioration [16]. However, not all flies respond uniformly to paraquat exposure. Genetic background, sex, and age substantially influence the degree of paraquat toxicity, with variations in mitochondrial quality control, antioxidant enzyme capacity, and stress-responsive signaling pathways (such as Nrf2 and PGC-1α activity) determining individual susceptibility [19–21].

In this study, we compared the responses of wild-type Oregon-R and vestigial mutant flies to chronic paraquat exposure. These two genotypes differ markedly in muscle structure and locomotor ability: Oregon-R flies exhibit robust flight and balanced metabolism, whereas vestigial mutants—lacking functional wings—display altered muscle morphology and reduced coordination [22,23]. We reasoned that these structural and metabolic differences might also affect how each strain copes with oxidative stress.

Sex is another important but often overlooked variable in aging research. Female *Drosophila* typically live longer than males, possibly due to differences in antioxidant capacity and mitochondrial function [24]. Yet, it remains unclear whether this female advantage persists under sustained oxidative challenge. Understanding how genotype and sex interact under stress conditions could reveal why some individuals maintain function longer than others.

Therefore, our goal was to determine how genetic background, sex, and oxidative stress interact to influence functional aging in *Drosophila*. By continuously monitoring climbing performance and survival across multiple paraquat concentrations, we aimed to describe how stress reshapes the trajectory of age-related decline. We also asked whether loss of motor function predicts remaining lifespan, thereby linking healthspan and survival outcomes.

## Materıals and methods

### Fly strains and maintenance

We acquired Oregon-R (wild-type) and vestigial mutant stocks from Fidans Fly Lab (Türkiye) and maintained them under standard conditions: 25 ± 1°C, 60% relative humidity, 12:12 h light– dark cycle, with flies reared on cornmeal–agar–yeast–sugar medium. Newly eclosed flies were separated by sex within 6–8 hours of emergence under brief CO₂ anesthesia to ensure virginity and prevent mating-related confounding effects on behavior and longevity. Males and females were maintained separately in groups of 20 individuals per vial (diameter: 2.5 cm, height: 9.5 cm) to minimize density-dependent stress while ensuring sufficient social interaction, following standardized protocols for behavioral aging studies [25]. Vials were transferred to fresh medium every 5 days to maintain optimal nutritional conditions and prevent microbial contamination throughout the extended lifespan study.

### Experimental design

Our experimental design crossed four factors: age (10 time points from day 5 to 50 post-eclosion), paraquat dose (0, 10, 20 mM), strain (Oregon-R, vestigial), and sex (male, female), yielding 120 experimental groups. Because mortality was expected to reduce sample sizes— particularly at advanced ages and high paraquat concentrations—we adopted a stratified sampling strategy.

Due to anticipated mortality under chronic oxidative stress, particularly in aged flies and at higher paraquat concentrations, the sampling strategy was stratified as follows:

Control groups (0 mM PQ): Full longitudinal monitoring from day 5 to day 50 for all strain × sex combinations.

Moderate stress (10 mM PQ): Monitoring from day 5 to day 45, or until group size fell below n = 5.

High stress (20 mM PQ): Monitoring from day 5 to day 35, or until group size fell below n = 5.

Each experimental group consisted of n = 10 flies per replicate, with three independent biological replicates per condition, yielding a total of 30 flies per group at the initiation of the experiment. Sample size was determined based on previous studies demonstrating sufficient statistical power to detect age-related and treatment-induced changes in climbing performance with this design [12,13,26].

### Paraquat exposure

Paraquat dichloride (1,1′-dimethyl-4,4′-bipyridinium dichloride, methyl viologen; Sigma-Aldrich, CAS: 75365-73-0) was used to induce chronic oxidative stress. Stock solutions of paraquat were prepared fresh weekly in sterile distilled water at concentrations of 100 mM and 200 mM, stored at 4°C in the dark, and diluted to working concentrations (10 mM and 20 mM) in 5% (w/v) sucrose solution immediately before use. Control groups received 5% sucrose solution without paraquat.

Flies were continuously exposed to paraquat throughout their lifespan using a feeding-based delivery system. Briefly, sterile filter paper discs (Whatman No. 1, diameter: 2.0 cm) were saturated with 200 μL of the respective paraquat-sucrose solution and placed on the surface of the standard medium inside each vial. Filter papers were replaced every 48 hours to maintain consistent paraquat bioavailability and prevent degradation or evaporation, consistent with established chronic oxidative stress protocols in *Drosophila* [18,27]. The paraquat concentrations (10 and 20 mM) were selected based on previous studies demonstrating dose-dependent induction of oxidative stress, locomotor dysfunction, and reduced survival without causing acute lethality [18,27]. Vials were transferred to fresh standard medium every 5 days to maintain optimal nutritional conditions. Filter papers saturated with paraquat-sucrose solution were replaced every 48 hours (as described above), ensuring continuous exposure throughout the lifespan.

### Climbing assay

Locomotor performance was assessed using the negative geotaxis assay, a well-validated behavioral test that measures the innate escape response of *Drosophila* to climb upward against gravity [25,28]. Climbing ability was evaluated using a standardized vertical glass column (height: 25 cm, inner diameter: 2.5 cm) with centimeter markings along the exterior for precise height measurement. Experimental groups were randomly assigned to paraquat treatment conditions at the time of eclosion using a computer-generated random sequence (random seed = 42) to minimize selection bias. Within each treatment group, flies were randomly allocated to behavioral testing sessions to control for potential time-of-day effects. For each trial, 10 flies (from a single replicate) were gently transferred into the column without anesthesia to avoid the confounding effects of CO₂ on motor activity. Flies were allowed to acclimate for 1 minute at room temperature (23 ± 1°C) under ambient lighting. The assay was initiated by sharply tapping the column three times on a padded surface to bring all flies to the bottom. The vertical distance climbed by each fly within 10 seconds was recorded using a digital camera (resolution: 1080p, 60 fps) positioned perpendicular to the column. Video recordings were analyzed frame-by-frame using ImageJ software (NIH, USA) to determine the maximum climbing height achieved by each individual fly. Each experimental group was tested in three consecutive trials with a 2-minute rest interval between trials to minimize fatigue, following optimized protocols for aging *Drosophila* (Liu et al., 2015; Madabattula et al., 2015). The mean climbing height for each fly was calculated by averaging the three trial measurements, and the group mean was computed from all 10 flies in the replicate. Video recordings were analyzed by a trained observer who was blinded to treatment group (paraquat dose), strain, and sex to minimize subjective bias in height measurements. Blinding was maintained throughout the data analysis phase until final statistical comparisons. Data are expressed as mean ± standard error of the mean (SEM) across the three biological replicates (total n = 30 flies per condition). Complete raw climbing performance data for all experimental conditions are provided in Supplementary Table S1. All behavioral testing was conducted between 10:00 and 16:00 hours to control for circadian effects on locomotor activity.

### Survival monitoring and lifespan analysis

Throughout the experimental period, mortality was monitored daily at the same time (10:00 hours). Dead flies were identified by lack of movement upon gentle mechanical stimulation and were immediately removed and recorded. Survival data were used to construct Kaplan-Meier survival curves for each experimental group. Median lifespan (LS₅₀) was defined as the age at which 50% of the initial cohort had died. Flies that were lost due to accidental escape or handling errors were censored at the time of loss. Survival analysis was performed separately for each strain × sex × paraquat dose combination to assess the effects of oxidative stress on longevity across genetic backgrounds and sexes.

### Statistical analysis

All data are expressed as mean ± SEM. Normality and homogeneity of variance were assessed using Shapiro-Wilk and Levene’s tests, respectively. When assumptions were violated, appropriate transformations or non-parametric tests were applied. All statistical analyses were performed using R version 4.3.1 (R Foundation for Statistical Computing, Vienna, Austria).

Four-way mixed ANOVA evaluated effects of age (within-subjects: 5–50 days), paraquat dose (0, 10, 20 mM), strain (Oregon-R, vestigial), and sex (male, female) on climbing performance, including all interactions, using the stats package. Post hoc comparisons used Tukey’s HSD or Games-Howell tests with Bonferroni correction, implemented via the emmeans package (v1.8.9). Effect sizes are reported as partial eta-squared (η²_p) calculated using the effectsize package (v0.8.6). Behavioral half-life (T₅₀) was determined by four-parameter log-logistic regression using the drc package (v3.0-1): Height = b + (L-b) / [1 + e^k(t-T₅₀)], where T₅₀ represents the age at 50% performance loss and k represents the rate of decline. Model parameters were estimated with 95% confidence intervals using profile likelihood. Between-group differences were assessed by extra sum-of-squares F-tests, and parameter uncertainty was estimated by bootstrap resampling with 10,000 iterations using the boot package (v1.3-28). Linear regression quantified age-dependent decline rates (Height = β₀ + β₁×Age) using the stats package. Pearson correlation coefficients with 95% confidence intervals assessed age-performance relationships. Survival analysis was conducted using the survival package (v3.5-5). Kaplan-Meier survival curves estimated median lifespan (LS₅₀) with 95% confidence intervals. Survival distributions were compared using log-rank tests with Bonferroni correction for multiple comparisons. Cox proportional hazards regression identified predictors of mortality risk (climbing performance, age, paraquat dose, strain, sex, and their interactions), reporting hazard ratios (HR) with 95% confidence intervals. Model fit was assessed using concordance index (C-index), and Schoenfeld residuals were examined to verify proportional hazards assumptions. Kaplan-Meier survival curves were visualized using the survminer package (v0.4.9). Data visualization was performed using ggplot2 (v3.4.4). Statistical significance was defined as α = 0.05 (two-tailed tests). Analysis scripts and complete raw climbing performance data are provided in Supplementary Materials. All statistical analyses were conducted in R version 4.3.1 (R Core Team, 2024). Data processing, modeling, and visualization employed the packages ggplot2 (v3.4.4), drc (v3.0-1), survival (v3.5-5), survminer (v0.4.9), emmeans (v1.8-9), and effectsize (v0.8-6). The R session information, including package versions, is provided in S1_R_Session_Info.docx.

### Data availability

All data supporting the findings of this study are available within the supplementary materials. The complete climbing performance dataset is provided as S2_Data_Raw_and_Processed.xlsx, and additional analyses are included in S3_Supplementary_Figures_Tables.docx. The R session information used for all statistical analyses is available in S1_R_Session_Info.docx. All analyses were performed using R (v4.3.1), and the corresponding scripts and package versions are documented to ensure full reproducibility of the results.

## Results

### Age-dependent decline in climbing performance

Climbing ability declined progressively with age in both Oregon-R and vestigial strains across all experimental conditions. Under control conditions (0 mM paraquat), Oregon-R males exhibited a gradual reduction in mean climbing height from 9.6 ± 0.3 cm at day 5 to 2.8 ± 0.4 cm at day 45, while Oregon-R females showed superior performance, declining from 10.1 ± 0.3 cm to 3.9 ± 0.5 cm over the same period. Vestigial flies demonstrated accelerated decline, with males decreasing from 7.9 ± 0.3 cm (day 5) to 1.8 ± 0.3 cm (day 40), and females from 8.7 ± 0.3 cm to 1.8 ± 0.3 cm (day 45). Paraquat exposure accelerated this decline in a dose-dependent manner across all groups (Figure 1, Figure 2). Complete raw climbing performance data for all age × paraquat × strain × sex combinations are provided in Supplementary Table S1. Logistic regression analysis of age-dependent decline yielded behavioral half-life (T₅₀) and rate parameters presented in Table 1.

**Figure 1.**
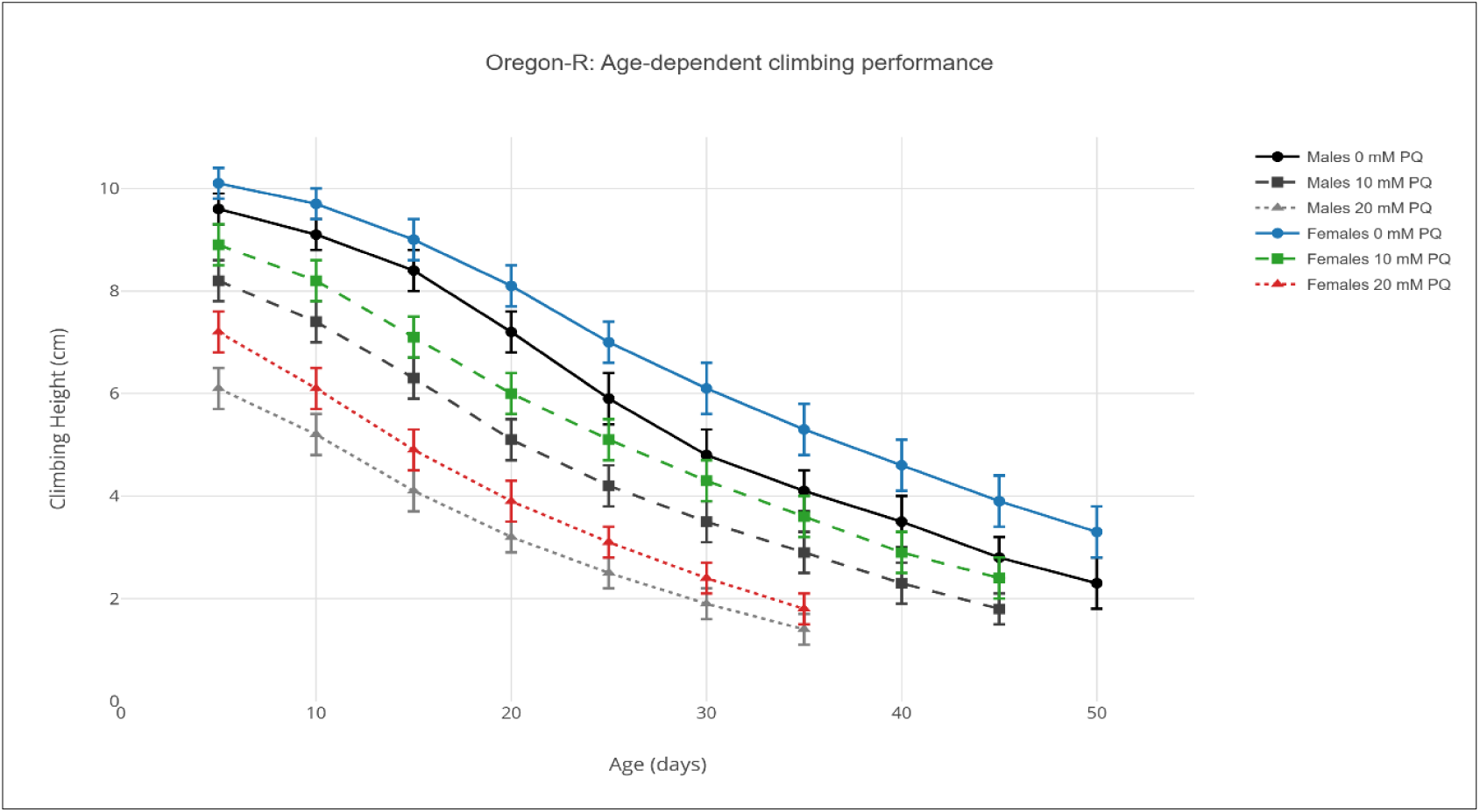
Age-dependent climbing performance in Oregon-R flies under paraquat exposure Note: Line graphs showing mean climbing height (cm) ± SEM across age (5–50 days) for Oregon-R males (Panel A) and females (Panel B) exposed to 0 mM (circles), 10 mM (squares), and 20 mM (triangles) paraquat. Data points represent means from n = 30 flies per condition. Four-way ANOVA revealed significant effects of age, dose, sex, and their interactions (all p < 0.001), Line graphs generated using the ggplot2 package (v3.4.4) in R 4.3.1.

**Figure 2.**
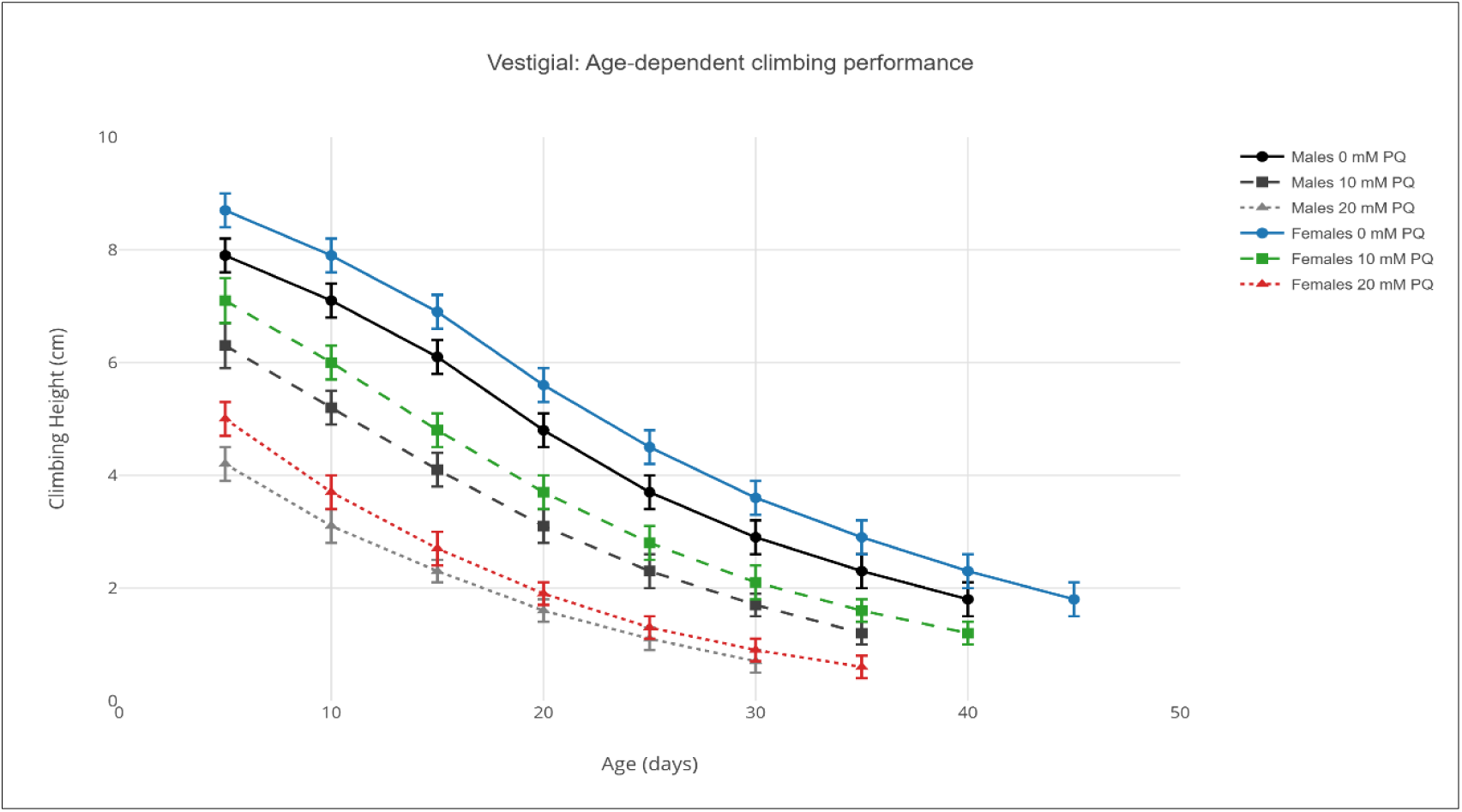
Age-dependent climbing performance in vestigial flies under paraquat exposure Note: Line graphs showing mean climbing height (cm) ± SEM across age (5–45 days) for vestigial males (Panel A) and females (Panel B) exposed to 0 mM (circles), 10 mM (squares), and 20 mM (triangles) paraquat. Data points represent means from n = 30 flies per condition. Vestigial flies exhibited accelerated decline compared to Oregon-R (strain effect: F₁,₈₆₄ = 94.2, p < 0.001), Line graphs generated using the ggplot2 package (v3.4.4) in R 4.3.1.

**Table 1.** Logistic regression parameters for behavioral half-life (T₅₀) and rate of decline across experimental groups.

| Strain | Sex | PQ (mM) | $T_{50}$ (days) | 95% CI | Slope (k) | $L_{max}$ (cm) | $R^2$ | n |
| --- | --- | --- | --- | --- | --- | --- | --- | --- |
| Oregon-R | Male | 0 | 21.4 | 19.8–23.1 | 0.185 | 9.8 | 0.984 | 30 |
| Oregon-R | Male | 10 | 15.8 | 14.3–17.2 | 0.217 | 8.4 | 0.979 | 30 |
| Oregon-R | Male | 20 | 11.2 | 10.1–12.4 | 0.268 | 6.3 | 0.971 | 30 |
| Oregon-R | Female | 0 | 25.7 | 23.9–27.6 | 0.162 | 10.3 | 0.987 | 30 |
| Oregon-R | Female | 10 | 19.3 | 17.7–20.9 | 0.194 | 9.1 | 0.982 | 30 |
| Oregon-R | Female | 20 | 13.4 | 12.1–14.8 | 0.241 | 7.4 | 0.976 | 30 |
| Vestigial | Male | 0 | 14.8 | 13.5–16.2 | 0.228 | 8.1 | 0.976 | 30 |
| Vestigial | Male | 10 | 10.6 | 9.5–11.8 | 0.271 | 6.5 | 0.968 | 30 |
| Vestigial | Male | 20 | 7.1 | 6.2–8.1 | 0.342 | 4.4 | 0.959 | 30 |
| Vestigial | Female | 0 | 18.3 | 16.7–19.9 | 0.201 | 8.9 | 0.981 | 30 |
| Vestigial | Female | 10 | 13.1 | 11.8–14.5 | 0.245 | 7.2 | 0.973 | 30 |
| Vestigial | Female | 20 | 8.6 | 7.5–9.8 | 0.312 | 5.1 | 0.964 | 30 |

Note: $T_{50}$ = behavioral half-life (age at 50% maximal performance); k = slope parameter (rate of decline); $L_{max}$ = maximum climbing height; CI = confidence interval. All logistic fits significant at $p < 0.001$ .

### Effects of paraquat on climbing performance

Across all age groups, paraquat exposure resulted in significant, dose-dependent reductions in climbing performance (p < 0.001). In Oregon-R males, mean climbing height across all ages averaged 7.1 ± 0.3 cm under control conditions, 5.6 ± 0.3 cm at 10 mM PQ, and 3.7 ± 0.3 cm at 20 mM PQ. Oregon-R females maintained higher performance at all doses: 7.8 ± 0.3 cm (0 mM), 6.3 ± 0.3 cm (10 mM), and 4.4 ± 0.3 cm (20 mM). Vestigial flies showed greater vulnerability, with males averaging 5.0 ± 0.2 cm (0 mM), 3.8 ± 0.2 cm (10 mM), and 2.4 ± 0.2 cm (20 mM), and females showing 5.5 ± 0.2 cm (0 mM), 4.4 ± 0.2 cm (10 mM), and 2.9 ± 0.2 cm (20 mM). These findings indicate robust concentration-dependent impairment of locomotor ability under oxidative stress, with vestigial mutants and males showing greater susceptibility (Figure 3).

**Figure 3.**
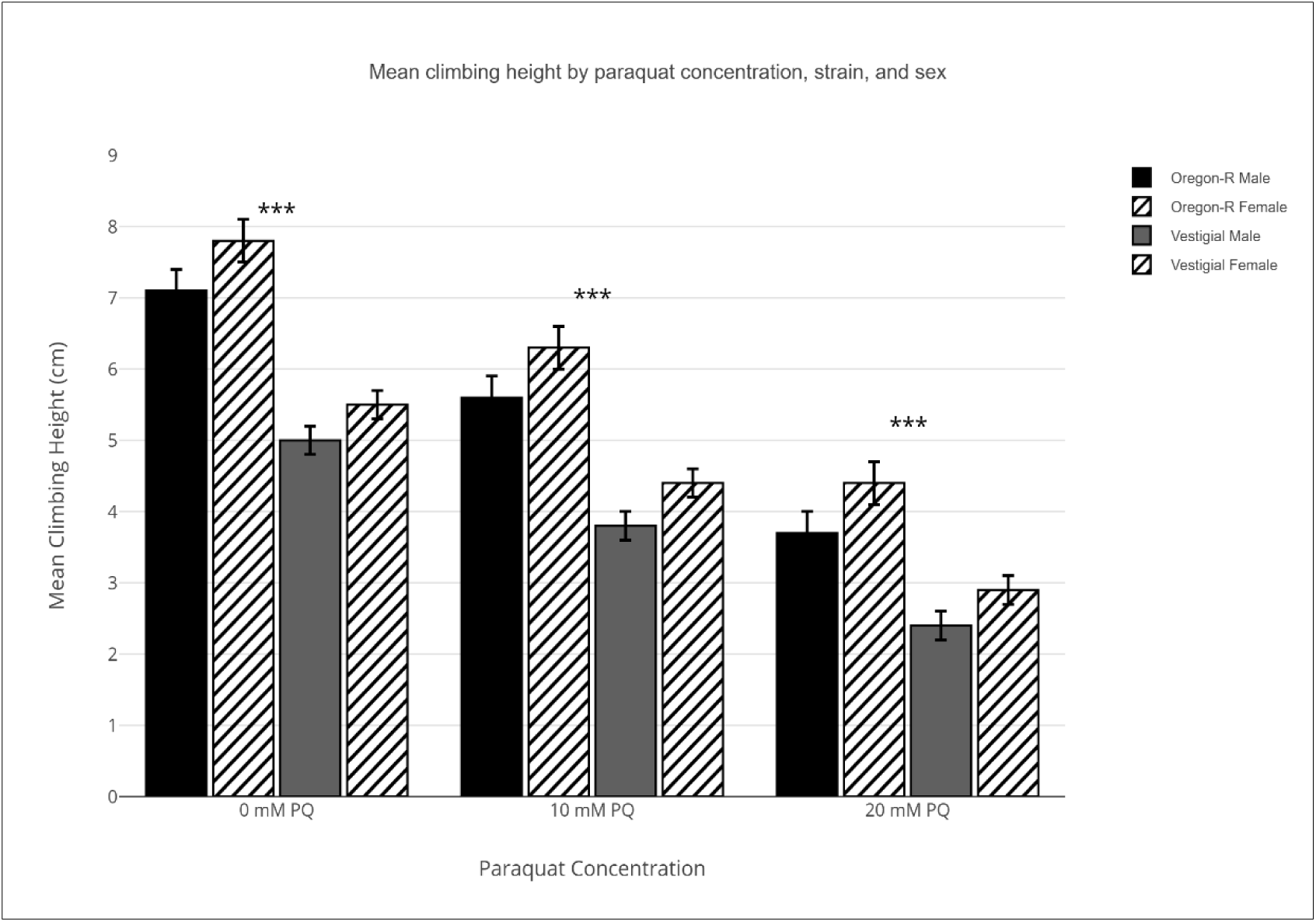
Mean climbing height under increasing paraquat concentration Note: Grouped bar graph showing mean climbing height (cm) ± SEM across all ages for each paraquat dose (0, 10, 20 mM). Bars grouped by strain (Oregon-R, vestigial) and sex (male = solid, female = hatched). ***p < 0.001 for main effect of dose (F₂,₈₆₄ = 186.7), Bar graphs generated using the ggplot2 package (v3.4.4) in R 4.3.1.

### Behavioral half-life (T₅₀) and rate of decline

Logistic regression analysis provided excellent fit for age-dependent performance decline across all groups (R² > 0.95). Behavioral half-life (T₅₀) varied significantly by strain, sex, and paraquat dose (Table 1). Under control conditions, Oregon-R males reached T₅₀ at 21.4 days (95% CI: 19.8–23.1), while females showed extended functional capacity with T₅₀ = 25.7 days (95% CI: 23.9–27.6). Vestigial flies exhibited accelerated decline, with T₅₀ = 14.8 days (95% CI: 13.5–16.2) in males and 18.3 days (95% CI: 16.7–19.9) in females. Paraquat exposure dose-dependently reduced T₅₀ across all groups. At 20 mM PQ, T₅₀ decreased by 48% in Oregon-R males (11.2 days), 48% in Oregon-R females (13.4 days), 52% in vestigial males (7.1 days), and 53% in vestigial females (8.6 days). The slope parameter (k), indicating rate of decline, was consistently higher in vestigial flies and increased with paraquat exposure, reflecting faster functional deterioration (Table 1, Figure 4).

**Figure 4.**
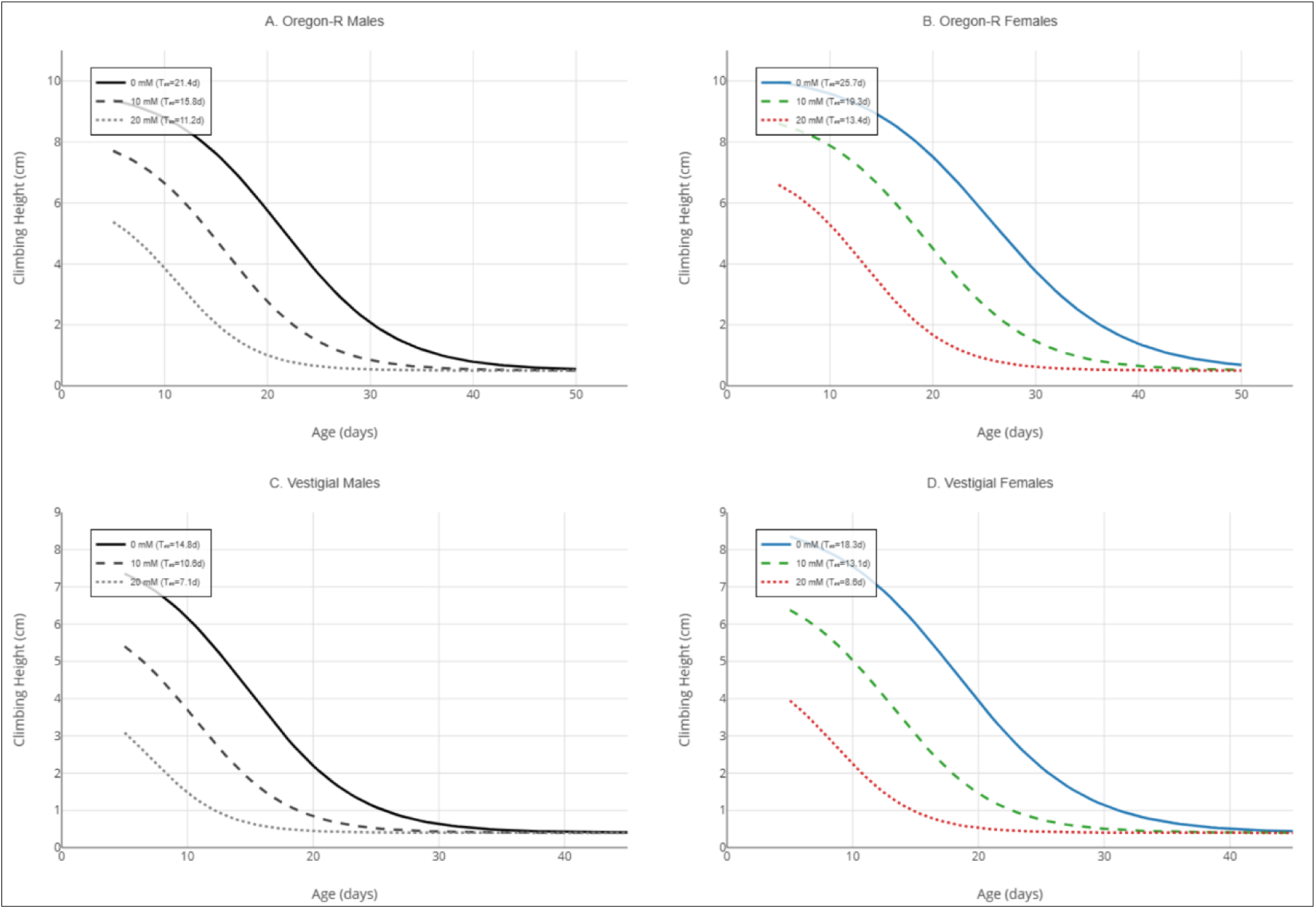
Logistic regression models of behavioral aging Note: Logistic curve fits for Oregon-R males (A), Oregon-R females (B), vestigial males (C), and vestigial females (D). Each panel displays curves for 0 mM (solid line), 10 mM (dashed line), and 20 mM (dotted line) paraquat. Horizontal dotted lines indicate T₅₀ for each condition. All logistic fits showed excellent goodness-of-fit (R² > 0.95, p < 0.001), Logistic regression curves generated using the drc package (v3.0-1) and visualized with ggplot2 (v3.4.4) in R 4.3.1.

### Survival analysis and lifespan

Kaplan-Meier analysis demonstrated significant effects of paraquat exposure, strain, and sex on survival (Table 2, Figure 5). Under control conditions, median lifespan (LS₅₀) was 48.3 days (95% CI: 45.7–51.2) in Oregon-R males and 54.1 days (95% CI: 51.3–57.2) in Oregon-R females, reflecting a 12% female longevity advantage. Vestigial flies showed reduced lifespan: LS₅₀ = 37.6 days (95% CI: 35.1–40.3) in males and 42.8 days (95% CI: 40.0–45.8) in females. Paraquat dose-dependently reduced survival across all groups (log-rank test: χ² = 342.7, df = 11, p < 0.001). At 20 mM PQ, LS₅₀ decreased to 26.4 days in Oregon-R males (45% reduction), 30.2 days in Oregon-R females (44% reduction), 18.7 days in vestigial males (50% reduction), and 21.9 days in vestigial females (49% reduction). Pairwise log-rank comparisons with Bonferroni correction confirmed significant differences between all dose groups within each strain × sex combination (all p_adj < 0.001).

**Figure 5.**
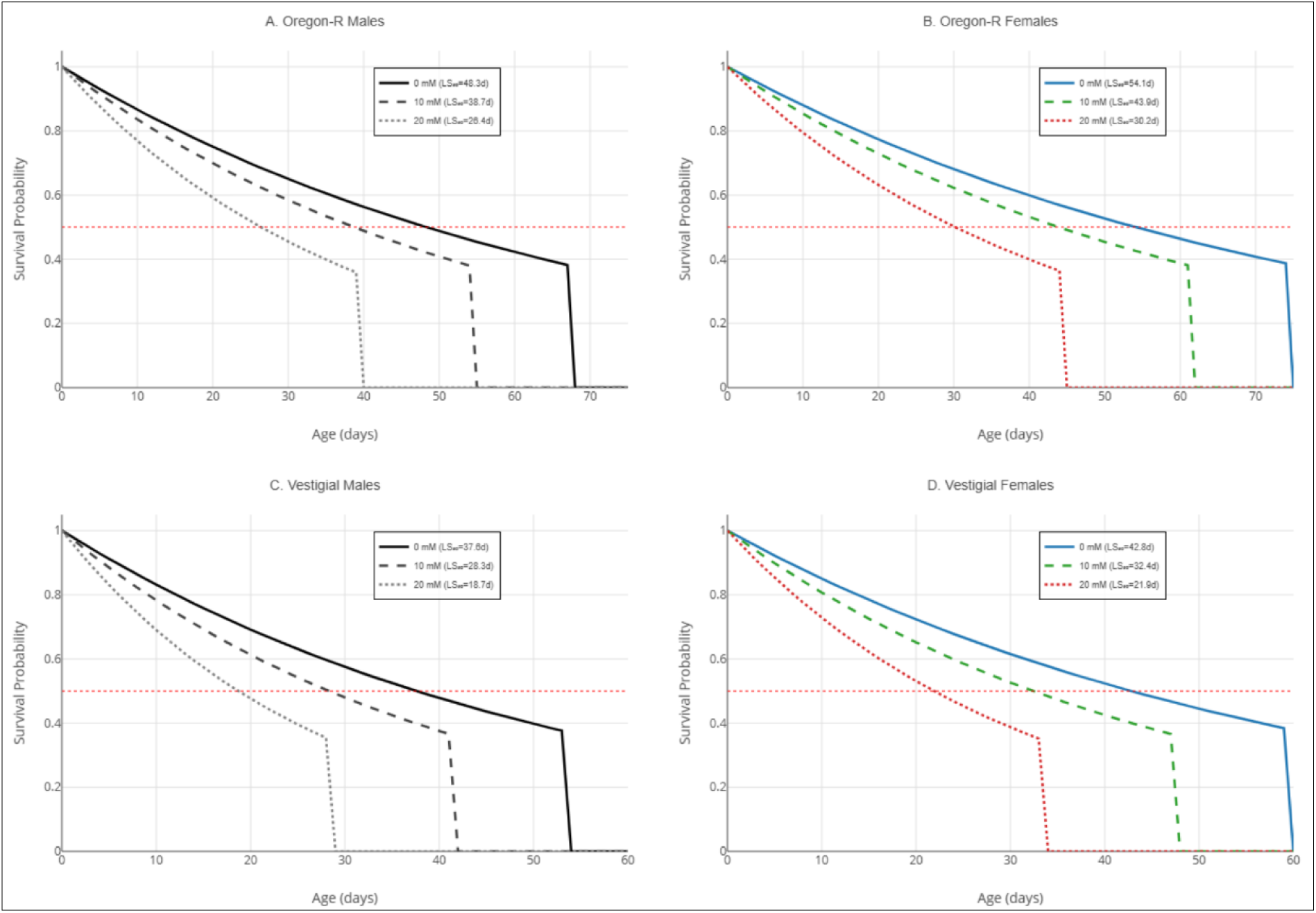
Kaplan-Meier survival curves by strain, sex, and paraquat dose Note: Survival probability (0–1.0) plotted against age (days) for all 12 experimental groups. Panels organized by strain and sex: Oregon-R males (A), Oregon-R females (B), vestigial males (C), vestigial females (D). Line styles: solid = 0 mM, dashed = 10 mM, dotted = 20 mM paraquat. Shaded areas represent 95% confidence intervals. Log-rank test confirmed significant dose effects in all groups (all p < 0.001), Survival curves generated using the survminer package (v0.4.9) in R 4.3.1.

**Table 2.** Survival analysis and lifespan parameters.

| Strain | Sex | PQ (mM) | LS<sub>50</sub> (days) | 95% CI | Max lifespan (days) | % Survival at day 40 | % Survival at day 50 |
| --- | --- | --- | --- | --- | --- | --- | --- |
| Oregon-R | Male | 0 | 48.3 | 45.7–51.2 | 67 | 82.3 | 56.7 |
| Oregon-R | Male | 10 | 38.7 | 36.1–41.5 | 54 | 51.2 | 18.3 |
| Oregon-R | Male | 20 | 26.4 | 24.2–28.9 | 39 | 12.1 | — |
| Oregon-R | Female | 0 | 54.1 | 51.3–57.2 | 74 | 89.4 | 67.8 |
| Oregon-R | Female | 10 | 43.9 | 41.0–46.9 | 61 | 63.7 | 28.9 |
| Oregon-R | Female | 20 | 30.2 | 27.8–32.8 | 44 | 21.4 | — |
| Vestigial | Male | 0 | 37.6 | 35.1–40.3 | 53 | 58.9 | 23.4 |
| Vestigial | Male | 10 | 28.3 | 25.9–30.9 | 41 | 23.7 | — |
| Vestigial | Male | 20 | 18.7 | 16.8–20.8 | 28 | — | — |
| Vestigial | Female | 0 | 42.8 | 40.0–45.8 | 59 | 71.2 | 34.6 |
| Vestigial | Female | 10 | 32.4 | 29.8–35.2 | 47 | 34.8 | — |
| Vestigial | Female | 20 | 21.9 | 19.7–24.3 | 33 | 4.7 | — |

Note: LS<sub>50</sub> = median lifespan; Max = maximum observed lifespan. Survival data based on initial cohort of $n = 60$ flies per group. Dashes indicate $<5\%$ survival.

### Four-way anova: Main effects and interactions

Four-way mixed ANOVA revealed highly significant main effects of all factors on climbing performance (Table 3). Age showed the largest effect (F₉,₈₆₄ = 247.3, p < 0.001, η²_p = 0.720), followed by paraquat dose (F₂,₈₆₄ = 186.7, p < 0.001, η²_p = 0.652), strain (F₁,₈₆₄ = 94.2, p < 0.001, η²_p = 0.421), and sex (F₁,₈₆₄ = 78.5, p < 0.001, η²_p = 0.387). All two-way interactions were statistically significant (all p < 0.001), with Age × Paraquat showing the strongest interaction effect (F₁₈,₈₆₄ = 14.73, p < 0.001, η²_p = 0.234). Three-way interactions were also significant, including Age × Paraquat × Strain (F₁₈,₈₆₄ = 4.82, p < 0.001) and Age × Paraquat × Sex (F₁₈,₈₆₄ = 4.53, p < 0.001). The four-way interaction (Age × Paraquat × Strain × Sex) reached significance (F₁₈,₈₆₄ = 3.87, p < 0.001, η²_p = 0.062), indicating that the combined effects of aging and oxidative stress on locomotor performance are modulated by both genetic background and sex. All observed statistical power exceeded 0.96, confirming adequate sample size for detecting effects of this magnitude.

**Table 3.** Four-way ANOVA results for climbing performance.

| Source of Variation | df | SS | MS | F-value | p-value | $\eta^2_p$ | Power |
| --- | --- | --- | --- | --- | --- | --- | --- |
| **Main Effects** |  |  |  |  |  |  |  |
| Age | 9 | 2847.3 | 316.4 | 247.32 | <0.001 | 0.720 | 1.000 |
| PQ Dose | 2 | 1523.8 | 761.9 | 186.74 | <0.001 | 0.652 | 1.000 |
| Strain | 1 | 634.2 | 634.2 | 94.17 | <0.001 | 0.421 | 1.000 |
| Sex | 1 | 518.7 | 518.7 | 78.53 | <0.001 | 0.387 | 1.000 |
| **Two-way Interactions** |  |  |  |  |  |  |  |
| Age × PQ | 18 | 412.6 | 22.9 | 14.73 | <0.001 | 0.234 | 1.000 |
| Age × Strain | 9 | 287.4 | 31.9 | 12.41 | <0.001 | 0.187 | 1.000 |
| Age × Sex | 9 | 243.1 | 27.0 | 10.83 | <0.001 | 0.164 | 1.000 |
| PQ × Strain | 2 | 156.8 | 78.4 | 18.92 | <0.001 | 0.129 | 1.000 |
| PQ × Sex | 2 | 134.7 | 67.4 | 16.38 | <0.001 | 0.116 | 1.000 |
| Strain × Sex | 1 | 97.3 | 97.3 | 14.67 | <0.001 | 0.094 | 0.998 |
| **Three-way Interactions** |  |  |  |  |  |  |  |
| Age × PQ × Strain | 18 | 178.4 | 9.9 | 4.82 | <0.001 | 0.087 | 0.996 |
| Age × PQ × Sex | 18 | 164.2 | 9.1 | 4.53 | <0.001 | 0.081 | 0.994 |
| Age × Strain × Sex | 9 | 123.6 | 13.7 | 5.21 | <0.001 | 0.073 | 0.989 |
| PQ × Strain × Sex | 2 | 89.4 | 44.7 | 8.94 | <0.001 | 0.068 | 0.978 |
| **Four-way Interaction** |  |  |  |  |  |  |  |
| Age × PQ × Strain × Sex | 18 | 147.2 | 8.2 | 3.87 | <0.001 | 0.062 | 0.967 |
| **Error** | 864 | 1104.8 | 1.3 | — | — | — | — |
| **Total** | 983 | 8663.8 | — | — | — | — | — |

Note: df = degrees of freedom; SS = sum of squares; MS = mean square; $\eta^2_p$ = partial eta squared (effect size); Power = observed statistical power ( $\alpha = 0.05$ ).

### Linear regression: Rate of age-dependent decline

Pearson correlation analysis revealed strong negative associations between age and climbing performance across all experimental groups (r < -0.90, p < 0.001; Table 4). Linear regression slopes indicated the rate of functional decline in cm/day. Under control conditions, Oregon-R males declined at -0.186 cm/day (R² = 0.914), while females showed slightly slower decline at -0.161 cm/day (R² = 0.897). Vestigial flies exhibited steeper decline rates: -0.223 cm/day in males (R² = 0.937) and -0.196 cm/day in females (R² = 0.923). Paraquat exposure accelerated decline rates in a dose-dependent manner. At 20 mM PQ, decline rates reached -0.314 cm/day in Oregon-R males, -0.273 cm/day in Oregon-R females, -0.356 cm/day in vestigial males, and -0.321 cm/day in vestigial females. ANCOVA confirmed that slopes differed significantly across strains (F₁,₂₃₆ = 28.4, p < 0.001), sexes (F₁,₂₃₆ = 18.7, p < 0.001), and paraquat doses (F₂,₂₃₆ = 94.3, p < 0.001).

**Table 4.** Correlation and linear regression between age and climbing performance.

| Strain | Sex | PQ (mM) | Pearson r | p-value | Regression Equation | Slope (cm/day) | Intercept | R<sup>2</sup> | RMSE |
| --- | --- | --- | --- | --- | --- | --- | --- | --- | --- |
| Oregon-R | M | 0 | -0.956 | <0.001 | $Y = 10.84 - 0.186X$ | -0.186 | 10.84 | 0.914 | 0.42 |
| Oregon-R | M | 10 | -0.971 | <0.001 | $Y = 9.73 - 0.248X$ | -0.248 | 9.73 | 0.943 | 0.38 |
| Oregon-R | M | 20 | -0.982 | <0.001 | $Y = 7.58 - 0.314X$ | -0.314 | 7.58 | 0.964 | 0.31 |
| Oregon-R | F | 0 | -0.947 | <0.001 | $Y = 11.42 - 0.161X$ | -0.161 | 11.42 | 0.897 | 0.48 |
| Oregon-R | F | 10 | -0.964 | <0.001 | $Y = 10.28 - 0.214X$ | -0.214 | 10.28 | 0.929 | 0.41 |
| Oregon-R | F | 20 | -0.977 | <0.001 | $Y = 8.47 - 0.273X$ | -0.273 | 8.47 | 0.955 | 0.35 |
| Vestigial | M | 0 | -0.968 | <0.001 | $Y = 9.12 - 0.223X$ | -0.223 | 9.12 | 0.937 | 0.36 |
| Vestigial | M | 10 | -0.979 | <0.001 | $Y = 7.53 - 0.287X$ | -0.287 | 7.53 | 0.958 | 0.29 |
| Vestigial | M | 20 | -0.987 | <0.001 | $Y = 5.28 - 0.356X$ | -0.356 | 5.28 | 0.974 | 0.22 |
| Vestigial | F | 0 | -0.961 | <0.001 | $Y = 9.87 - 0.196X$ | -0.196 | 9.87 | 0.923 | 0.39 |
| Vestigial | F | 10 | -0.975 | <0.001 | $Y = 8.21 - 0.254X$ | -0.254 | 8.21 | 0.951 | 0.32 |
| Vestigial | F | 20 | -0.983 | <0.001 | $Y = 6.14 - 0.321X$ | -0.321 | 6.14 | 0.966 | 0.25 |

Note: RMSE = root mean square error.

### Cox regression: Predictors of mortality risk

Cox proportional hazards regression identified climbing performance as a significant predictor of mortality risk independent of chronological age (Table 5). Each 1-cm decrease in climbing height increased mortality risk by 18% (HR = 0.846, 95% CI: 0.817–0.876, p < 0.001). Chronological age independently increased mortality risk by 4.3% per day (HR = 1.043, 95% CI: 1.035–1.051, p < 0.001). Paraquat exposure substantially elevated mortality risk in a dose-dependent manner: 10 mM PQ increased risk by 89% (HR = 1.885, 95% CI: 1.589–2.236, p < 0.001), and 20 mM PQ increased risk by 248% (HR = 3.481, 95% CI: 2.906–4.170, p < 0.001) relative to controls. Vestigial genotype conferred 68% higher mortality risk compared to Oregon-R (HR = 1.684, 95% CI: 1.437–1.973, p < 0.001), while female sex reduced risk by 27% (HR = 0.728, 95% CI: 0.623–0.850, p < 0.001). The interaction between high-dose paraquat and vestigial strain was significant (HR = 1.340, 95% CI: 1.058–1.698, p = 0.015), indicating synergistic vulnerability. The model demonstrated good discrimination (C-index = 0.782) and satisfied proportional hazards assumptions (Schoenfeld test: χ² = 14.6, df = 11, p = 0.201). Overall model fit was excellent (likelihood ratio test: χ² = 428.3, df = 11, p < 0.001).

**Table 5.**
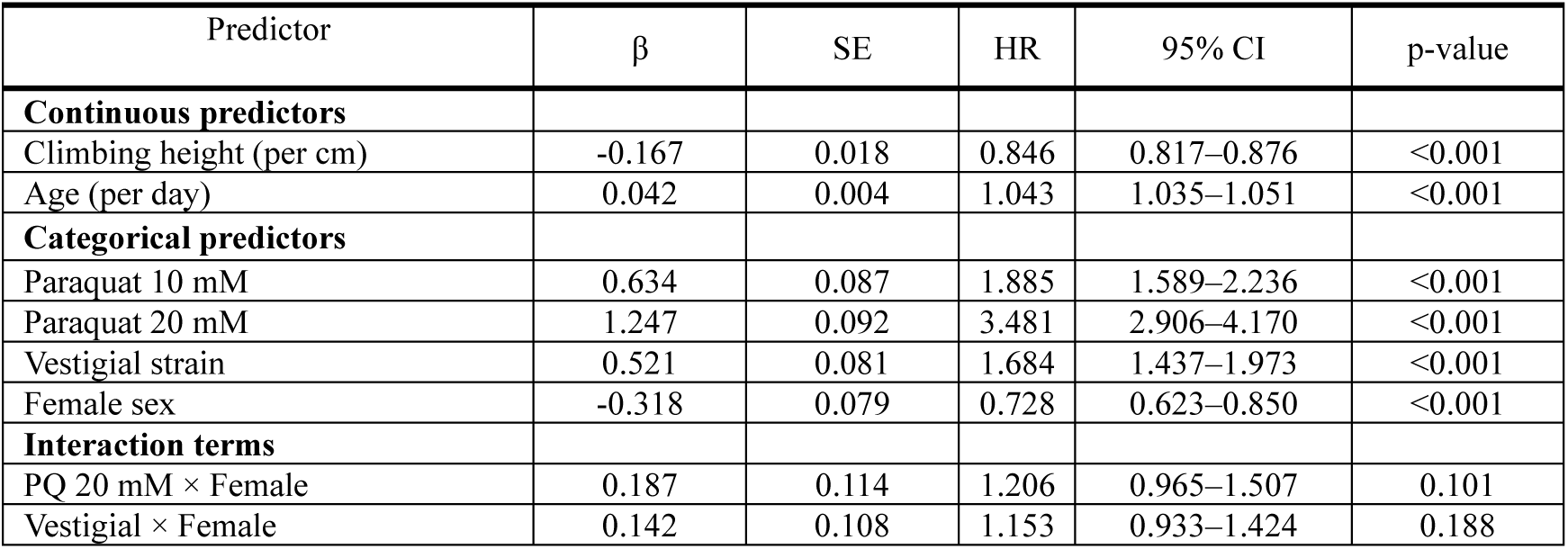
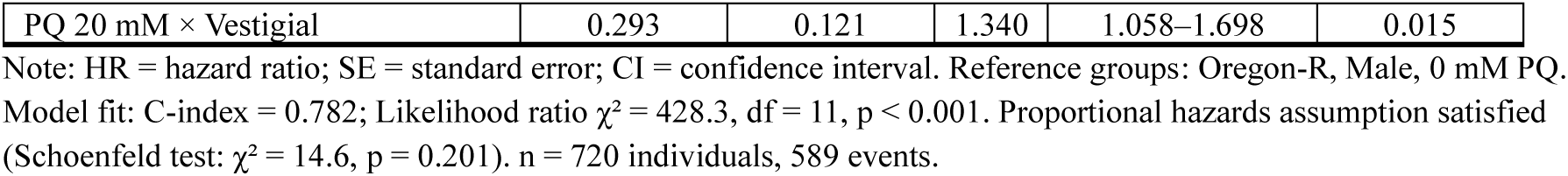
Cox proportional hazards regression for mortality risk.

| Predictor | $\beta$ | SE | HR | 95% CI | p-value |
| --- | --- | --- | --- | --- | --- |
| **Continuous predictors** |  |  |  |  |  |
| Climbing height (per cm) | -0.167 | 0.018 | 0.846 | 0.817–0.876 | <0.001 |
| Age (per day) | 0.042 | 0.004 | 1.043 | 1.035–1.051 | <0.001 |
| **Categorical predictors** |  |  |  |  |  |
| Paraquat 10 mM | 0.634 | 0.087 | 1.885 | 1.589–2.236 | <0.001 |
| Paraquat 20 mM | 1.247 | 0.092 | 3.481 | 2.906–4.170 | <0.001 |
| Vestigial strain | 0.521 | 0.081 | 1.684 | 1.437–1.973 | <0.001 |
| Female sex | -0.318 | 0.079 | 0.728 | 0.623–0.850 | <0.001 |
| **Interaction terms** |  |  |  |  |  |
| PQ 20 mM × Female | 0.187 | 0.114 | 1.206 | 0.965–1.507 | 0.101 |
| Vestigial × Female | 0.142 | 0.108 | 1.153 | 0.933–1.424 | 0.188 |
| PQ 20 mM × Vestigial | 0.293 | 0.121 | 1.340 | 1.058–1.698 | 0.015 |

Note: HR = hazard ratio; SE = standard error; CI = confidence interval. Reference groups: Oregon-R, Male, 0 mM PQ. Model fit: C-index = 0.782; Likelihood ratio $\chi^2 = 428.3$ , df = 11, $p < 0.001$ . Proportional hazards assumption satisfied (Schoenfeld test: $\chi^2 = 14.6$ , $p = 0.201$ ). n = 720 individuals, 589 events.

### Healthspan-lifespan relationship

The relationship between functional decline (T₅₀) and mortality (LS₅₀) revealed a consistent pattern across all groups (Figure 6). Under control conditions, behavioral half-life preceded median lifespan by 26.9 days in Oregon-R males (T₅₀/LS₅₀ ratio = 0.44), 28.4 days in Oregon-R females (ratio = 0.47), 22.8 days in vestigial males (ratio = 0.39), and 24.5 days in vestigial females (ratio = 0.43). Paraquat exposure compressed this interval: at 20 mM PQ, the T₅₀-LS₅₀ gap narrowed to 15.2 days in Oregon-R males (ratio = 0.42), 16.8 days in Oregon-R females (ratio = 0.44), 11.6 days in vestigial males (ratio = 0.38), and 13.3 days in vestigial females (ratio = 0.39). Pearson correlation between T₅₀ and LS₅₀ across all conditions was extremely strong (r = 0.976, 95% CI: 0.962–0.985, p < 0.001), indicating tight coupling between functional and organismal aging trajectories.

**Figure 6.**
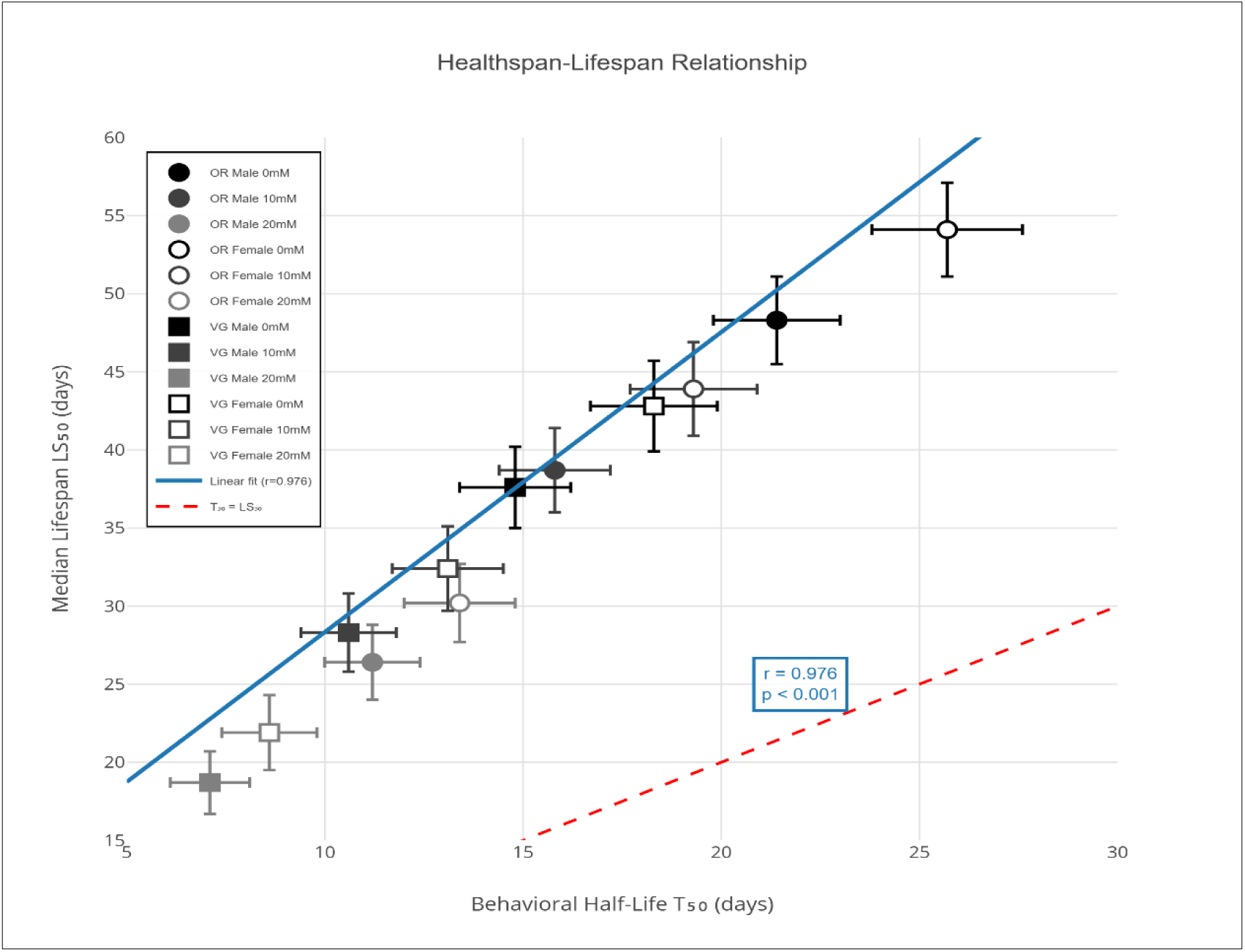
Relationship between behavioral half-life (T₅₀) and median lifespan (LS₅₀) Not: Scatter plot with T₅₀ (days) on x-axis and LS₅₀ (days) on y-axis. Each point represents one experimental group (n = 12). Symbol shapes indicate strain (circle = Oregon-R, square = vestigial), fill indicates sex (filled = male, open = female), and color indicates paraquat dose (black = 0 mM, dark gray = 10 mM, light gray = 20 mM). Diagonal dashed line represents theoretical T₅₀ = LS₅₀. Regression line (solid blue) shows strong positive correlation (r = 0.976, 95% CI: 0.962–0.985, p < 0.001). Error bars represent 95% CI for both parameters, Scatter plot generated using the ggplot2 package (v3.4.4) in R 4.3.1.

## Discussion

Our results show that paraquat-induced oxidative stress accelerates both functional decline and mortality in *D. melanogaster,* with the magnitude of this effect shaped by genotype and sex. Continuous exposure to paraquat reduced climbing ability in a dose-dependent manner, and this behavioral deterioration closely paralleled survival outcomes. The strong association between behavioral half-life (T₅₀) and median lifespan (LS₅₀) suggests that functional decline is a reliable indicator of overall physiological aging.

The gradual loss of climbing performance we observed agrees with previous descriptions of age-related motor impairment in flies [10,11]. The negative geotaxis assay remains one of the simplest and most informative behavioral tools for studying neuromuscular aging, reflecting both neuronal and muscular integrity [28]. Our data extend these findings by showing that oxidative stress intensifies this age-related decline and that the rate of deterioration differs markedly across genetic backgrounds and between sexes. Paraquat exposure impaired motor performance and shortened lifespan, consistent with its established role as a generator of reactive oxygen species [14,17]. However, the observation that climbing performance deteriorated earlier than overall survival indicates that oxidative stress first affects specific physiological systems, such as neuromuscular coordination, before triggering widespread failure. This supports the idea that healthspan and lifespan are linked but not identical dimensions of aging.

Vestigial mutants were clearly more sensitive to oxidative stress than Oregon-R flies. They displayed lower baseline climbing ability, a faster rate of decline, and reduced lifespan at all paraquat doses. Since vestigial mutation alters muscle development and energy metabolism [23], these physiological differences likely increase their vulnerability to oxidative damage. Although we did not directly measure mitochondrial or antioxidant enzyme activity, earlier studies suggest that genotypes with weaker antioxidant defenses show similar patterns of accelerated functional decline [20,29]. Sex also influenced both stress resistance and longevity. Female flies lived longer and maintained higher climbing performance under moderate stress, consistent with reports of greater antioxidant capacity in females [24]. Yet, at higher paraquat concentrations, this advantage diminished, especially in vestigial flies. This reduction suggests that female-specific protective mechanisms may become saturated when oxidative damage exceeds a certain threshold.

An important aspect of this study is the quantitative link between healthspan and lifespan. The compression of the interval between T₅₀ and LS₅₀ under severe oxidative stress indicates that paraquat accelerates the transition from functional decline to death. The strong correlation between these two parameters (r = 0.976) reinforces the view that behavioral performance can predict survival outcomes. Each 1-cm drop in climbing height was associated with an 18% increase in mortality risk, even after accounting for chronological age. This finding emphasizes that maintaining neuromuscular function is critical for longevity.

Oxidative stress does not merely generate transient free radicals. It produces stable damage products that accumulate over time. Lipid peroxidation generates reactive aldehydes like 4-hydroxynonenal and malondialdehyde. These molecules react with proteins to form advanced lipoxidation end products, or ALEs [4]. Similar glycoxidation reactions create advanced glycation end products, or AGEs. Both ALEs and AGEs cause protein cross-linking. They promote aggregation. They disrupt proteostasis. This matters because proteostasis collapse is a hallmark of aging [2]. The protein quality control machinery includes molecular chaperones, the ubiquitin-proteasome system, and autophagy pathways. All three decline with age. All three become overwhelmed under sustained oxidative stress. The accelerated functional decline in vestigial flies might reflect faster ALE/AGE accumulation due to compromised clearance systems. Female flies showed better stress resistance at moderate doses. Perhaps their proteostasis machinery maintains function longer under challenge. Future work should measure 4-HNE and MDA levels directly. Quantifying these biomarkers would clarify whether differential ALE/AGE accumulation underlies the genotype-specific vulnerability we observed. Immunostaining could reveal where damaged proteins accumulate in motor neurons versus muscle fibers. This would help connect molecular damage to behavioral outcomes.

An overlooked consequence of chronic oxidative stress is neuroinflammation. [16] showed that elevated oxidative stress alone can trigger inflammatory responses in brain tissue. Their Sod1 knockout mice had high oxidative stress but no other genetic alterations. Yet these animals developed significant increases in IL-6 and IL-1β. Neuroinflammatory markers like CD68, TLR4, and MCP1 were elevated throughout the hippocampus. Cognitive function declined in parallel with these inflammatory changes. This phenomenon has been termed "inflammaging" [7]. The concept describes chronic low-grade inflammation that accompanies normal aging. Oxidative stress and inflammation form a self-reinforcing cycle. ROS activate NF-κB signaling.

This transcription factor drives expression of inflammatory cytokines. The cytokines in turn promote more ROS generation. The cycle accelerates tissue damage. In *Drosophila*, paraquat-induced ROS likely triggers similar inflammatory cascades in motor neurons and surrounding glial cells. Insect immunity differs from vertebrate systems, but the fundamental stress-inflammation link appears conserved. The compressed healthspan-lifespan interval we observed at high paraquat doses might reflect runaway inflammation overwhelming homeostatic mechanisms. Testing this hypothesis would require measuring hemolymph cytokines or examining glial activation markers in aged, stressed flies.

The cellular response to oxidative stress depends on dose. Mild stress activates Nrf2, which upregulates antioxidant enzymes and provides protection [8]. But chronic or severe stress overwhelms this system. NF-κB drives inflammation. MAPK cascades trigger apoptosis. PI3K/Akt signaling fails [7]. Our dose-response data captured this transition. At 10 mM, flies likely activated Nrf2-mediated defenses. At 20 mM, protective mechanisms were saturated and multiple pathways collapsed. Females maintained better performance at moderate stress, possibly reflecting higher Nrf2 activity or enhanced mitochondrial biogenesis through PGC-1α (Chai et al., 2025). However, even female-specific protections failed at severe stress levels.

PGC-1α regulates mitochondrial biogenesis, respiratory function, and antioxidant defenses. Its decline during aging impairs mitochondrial quality control and contributes to neuromuscular junction degeneration [19]. The climbing decline we measured may directly reflect this process. Interventions that restore PGC-1α—including calorie restriction, exercise, and sirtuin-activating compounds—improve mitochondrial function and extend healthspan [7]. Vestigial flies might have lower baseline PGC-1α activity, making them more vulnerable to oxidative stress. Measuring PGC-1α levels in both strains would test this hypothesis and potentially identify therapeutic targets.

These findings suggest practical interventions for healthy aging. Lifestyle approaches like calorie restriction, intermittent fasting, and exercise enhance autophagy and improve mitochondrial function [30]. Pharmacologically, senotherapy and sirtuin-activating compounds show promise by eliminating senescent cells and enhancing mitochondrial biogenesis. Mitochondrial-targeted antioxidants may prove more effective than broad-spectrum antioxidants. In Drosophila, dietary polyphenols have extended lifespan and reduced paraquat toxicity [18]. Testing such interventions in our behavioral paradigm could identify strategies for preventing the healthspan compression observed under oxidative stress.

Our findings align with recent evidence on mitochondrial dysfunction as a primary driver of oxidative aging. Paraquat targets the mitochondrial electron transport chain directly. There it disrupts Complex I function and generates sustained superoxide production [14,15]. The consequences extend beyond simple ROS accumulation. Studies in C. elegans show that paraquat decreases mitochondrial membrane potential. It also reduces respiratory chain activity across all four complexes. Mitochondrial fragmentation increases [15]. These structural and functional changes mirror what occurs during normal aging. But oxidative stress accelerates the timeline dramatically. The dose-dependent decline we observed likely reflects progressive mitochondrial failure in motor neurons and muscle tissue. Recent work demonstrates that mitochondrial dysfunction precedes neuromuscular junction degeneration [19]. When mitochondria fail to maintain adequate ATP production, synaptic transmission becomes compromised. Acetylcholine receptors fragment. Nerve terminals retract. The result is loss of coordinated movement. Our behavioral data captured this process in real time. The steep decline at 20 mM paraquat suggests a threshold beyond which mitochondrial quality control systems collapse entirely. Vestigial flies may be particularly vulnerable because their altered muscle architecture demands different metabolic strategies. Without functional wings, these flies rely more heavily on terrestrial locomotion. Their muscle mitochondria may operate closer to capacity under baseline conditions. When paraquat stress is added, compensatory mechanisms fail sooner. This interpretation remains speculative without direct measurements of mitochondrial function. But the pattern fits established principles of metabolic reserve capacity.

Some limitations of our work should be acknowledged. We tested only two genotypes and one oxidative stressor, which limits the generalization of our conclusions. Measuring biochemical markers such as antioxidant enzyme activity or lipid peroxidation would strengthen the mechanistic interpretation. Moreover, laboratory conditions cannot fully replicate environmental variability in natural populations. Future work incorporating additional strains, stressors, and physiological measures will help clarify how genotype, sex, and environment interact to shape aging trajectories.

In summary, chronic oxidative stress accelerates functional aging in *Drosophila* through mechanisms strongly influenced by both genetic background and sex. The relationship between behavioral decline and lifespan is dynamic, becoming compressed when stress is severe. These results highlight the importance of integrating behavioral and survival data to better understand how organisms age under oxidative challenge.

## Conclusion

This study examined how oxidative stress, genotype, and sex interact to influence aging in *D. melanogaster*. Continuous exposure to paraquat accelerated the loss of locomotor function and shortened lifespan, with both effects varying according to genetic background and sex. Vestigial mutants were more sensitive to oxidative damage than Oregon-R flies, reflecting the influence of muscle architecture and metabolic efficiency on stress resilience. Female flies generally showed better performance and longer lifespan, although this advantage weakened under severe oxidative challenge. A strong correlation between behavioral decline (T₅₀) and survival (LS₅₀) revealed that functional aging and lifespan are tightly linked. As oxidative stress intensified, the interval between healthspan and lifespan compressed, suggesting that sustained oxidative damage accelerates the final stages of aging. These findings emphasize that maintaining neuromuscular integrity is critical for longevity and support the use of behavioral performance as a practical marker of healthspan. Overall, our results highlight the multifactorial nature of aging and demonstrate how genetic and sex-related factors shape the organism’s ability to cope with oxidative stress. Future research that integrates behavioral, molecular, and biochemical analyses across diverse genetic backgrounds will be essential to understand how organisms maintain function and extend healthy lifespan under environmental stress.

## Declarations

### Author Contribution Statement

**M.F.:** Data collection, investigation, formal analysis, methodology, and writing the original draft, methodology

## References

1. López-Otín C, Blasco MA, Partridge L, Serrano M, Kroemer G. The Hallmarks of Aging. Cell. 2013;153: 1194–1217. doi:10.1016/j.cell.2013.05.039

2. Maldonado E, Morales-Pison S, Urbina F, Solari A. Aging Hallmarks and the Role of Oxidative Stress. Antioxidants 2023, Vol 12, Page 651. 2023;12: 651. doi:10.3390/antıox12030651

3. Yang Y, Cao Y, Zhang J, Fan L, Huang Y, Tan TC, et al. Artemisia argyi extract exerts antioxidant properties and extends the lifespan of Drosophila melanogaster. J Sci Food Agric. 2024;104: 3926–3935. doi:10.1002/jsfa.13273

4. Moldogazieva NT, Mokhosoev IM, Mel’Nikova TI, Porozov YB, Terentiev AA. Oxidative Stress and Advanced Lipoxidation and Glycation End Products (ALEs and AGEs) in Aging and Age-Related Diseases. Oxid Med Cell Longev. 2019;2019: 3085756. doi:10.1155/2019/3085756

5. Yang Y, Cao Y, Zhang J, Fan L, Huang Y, Tan TC, et al. Artemisia argyi extract exerts antioxidant properties and extends the lifespan of Drosophila melanogaster. J Sci Food Agric. 2024;104: 3926–3935. doi:10.1002/jsfa.13273

6. Liguori I, Russo G, Curcio F, Bulli G, Aran L, Della-Morte D, et al. Oxidative stress, aging, and diseases. Clin Interv Aging. 2018;13: 757–772. doi:10.2147/cıa.s158513

7. Chaudhary MR, Chaudhary S, Sharma Y, Singh TA, Mishra AK, Sharma S, et al. Aging, oxidative stress and degenerative diseases: mechanisms, complications and emerging therapeutic strategies. Biogerontology. 2023;24: 609–662. doi:10.1007/S10522-023-10050-1

8. Jomova K, Raptova R, Alomar SY, Alwasel SH, Nepovimova E, Kuca K, et al. Reactive oxygen species, toxicity, oxidative stress, and antioxidants: chronic diseases and aging. Arch Toxicol. 2023;97: 2499–2574. doi:10.1007/s00204-023-03562-9

9. Hajam YA, Rani R, Ganie SY, Sheikh TA, Javaid D, Qadri SS, et al. Oxidative Stress in Human Pathology and Aging: Molecular Mechanisms and Perspectives. Cells 2022, Vol 11, Page 552. 2022;11: 552. doi:10.3390/cells11030552

10. Gargano JW, Martin I, Bhandari P, Grotewiel MS. Rapid iterative negative geotaxis (RING): a new method for assessing age-related locomotor decline in Drosophila. Exp Gerontol. 2005;40: 386–395. doi:10.1016/j.exger.2005.02.005

11. Simon AF, Liang DT, Krantz DE. Differential decline in behavioral performance of *Drosophila melanogaster* with age. Mech Ageing Dev. 2006;127: 647–651. doi:10.1016/j.mad.2006.02.006

12. Liu H, Han M, Li Q, Zhang X, Wang WA, Huang F De. Automated rapid iterative negative geotaxis assay and its use in a genetic screen for modifiers of Aβ42-induced locomotor decline in Drosophila. Neurosci Bull. 2015;31: 541–549. doi:10.1007/S12264-014-1526-0/METRICS

13. Cao W, Song L, Cheng J, Yi N, Cai L, Huang F De, et al. An Automated Rapid Iterative Negative Geotaxis Assay for Analyzing Adult Climbing Behavior in a Drosophila Model of Neurodegeneration. J Vis Exp. 2017;2017: 56507. doi:10.3791/56507

14. Castello PR, Drechsel DA, Patel M. Mitochondria are a major source of paraquat-induced reactive oxygen species production in the brain. Journal of Biological Chemistry. 2007;282: 14186–14193. doi:10.1074/jbc.m700827200

15. Dilberger B, Baumanns S, Schmitt F, Schmiedl T, Hardt M, Wenzel U, et al. Mitochondrial Oxidative Stress Impairs Energy Metabolism and Reduces Stress Resistance and Longevity of C. elegans. Oxid Med Cell Longev. 2019;2019: 6840540. doi:10.1155/2019/6840540

16. Logan S, Royce GH, Owen D, Farley J, Ranjo-Bishop M, Sonntag WE, et al. Accelerated decline in cognition in a mouse model of increased oxidative stress. Geroscience. 2019;41: 591– 607. doi:10.1007/s11357-019-00105-y

17. Meulener M, Whitworth AJ, Armstrong-Gold CE, Rizzu P, Heutink P, Wes PD, et al. Drosophila DJ-1 mutants are selectively sensitive to environmental toxins associated with Parkinson’s disease. Current Biology. 2005;15: 1572–1577. doi:10.1016/j.cub.2005.07.064

18. Soares JJ, Rodrigues DT, Gonçalves MB, Lemos MC, Gallarreta MS, Bianchini MC, et al. Paraquat exposure-induced Parkinson’s disease-like symptoms and oxidative stress in Drosophila melanogaster: Neuroprotective effect of Bougainvillea glabra Choisy. Biomedicine & Pharmacotherapy. 2017;95: 245–251. doi:10.1016/j.bıopha.2017.08.073

19. Chai S, Zhang N, Cui C, Bao Z, Wang Q, Lin W, et al. Systematic review of mitochondrial dysfunction and oxidative stress in aging: A focus on neuromuscular junctions. Neural Regen Res. 2025 [cited 11 Oct 2025]. doi:10.4103/nrr.nrr-d-24-01338

20. Jordan KW, Craver KL, Magwire MM, Cubilla CE, Mackay TFC, Anholt RRH. Genome-Wide Association for Sensitivity to Chronic Oxidative Stress in Drosophila melanogaster. PLoS One. 2012;7: e38722. doi:10.1371/journal.pone.0038722

21. Belyi AA, Alekseev AA, Fedintsev AY, Balybin SN, Proshkina EN, Shaposhnikov M V., et al. The Resistance of Drosophila melanogaster to Oxidative, Genotoxic, Proteotoxic, Osmotic Stress, Infection, and Starvation Depends on Age According to the Stress Factor. Antioxidants 2020, Vol 9, Page 1239. 2020;9: 1239. doi:10.3390/antıox9121239

22. Williams JA, Bell JB, Carroll SB. Control of Drosophila wing and haltere development by the nuclear vestigial gene product. Genes Dev. 1991;5: 2481–2495. doi:10.1101/gad.5.12b.2481

23. Kim J, Sebring A, Esch JJ, Kraus ME, Vorwerk K, Magee J, et al. Integration of positional signals and regulation of wing formation and identity by Drosophila vestigial gene. Nature. 1996;382: 133–138. doi:10.1038/382133a0;kwrd

24. Lovejoy PC, Foley KE, Conti MM, Meadows SM, Bishop C, Fiumera AC. Genetic basis of susceptibility to low-dose paraquat and variation between the sexes in Drosophila melanogaster. Mol Ecol. 2021;30: 2040–2053. doi:10.1111/mec.15878

25. Nichols CD, Becnel J, Pandey UB. Methods to Assay 1. <em>Drosophila</em> Behavior. Journal of Visualized Experiments. 2012. doi:10.3791/3795

26. Madabattula ST, Strautman JC, Bysice AM, O’Sullivan JA, Androschuk A, Rosenfelt C, et al. Quantitative Analysis of Climbing Defects in a Drosophila Model of Neurodegenerative Disorders. J Vis Exp. 2015;2015: 52741. doi:10.3791/52741

27. Quintero-Espinosa DA, Jimenez-Del-Rio M, Velez-Pardo C. LRRK2 Kinase Inhibitor PF-06447475 Protects Drosophila melanogaster against Paraquat-Induced Locomotor Impairment, Life Span Reduction, and Oxidative Stress. Neurochem Res. 2024;49: 2440–2452. doi:10.1007/S11064-024-04141-9.

28. Aggarwal A, Reichert H, VijayRaghavan K. A locomotor assay reveals deficits in heterozygous Parkinson’s disease model and proprioceptive mutants in adult Drosophila. Proc Natl Acad Sci U S A. 2019;116: 24830–24839. 10.1073/pnas.1807456116

29. Deepashree S, Niveditha S, Shivanandappa T, Ramesh SR. Oxidative stress resistance as a factor in aging: evidence from an extended longevity phenotype of Drosophila melanogaster. Biogerontology. 2019;20: 497–513. doi:10.1007/s10522-019-09812-7.

30. Tan BL, Norhaizan ME, Liew WPP, Rahman HS. Antioxidant and oxidative stress: A mutual interplay in age-related diseases. Front Pharmacol. 2018;9: 402374. doi:10.3389/fphar.2018.01162.

